# DO-MS: Data-Driven Optimization of Mass Spectrometry Methods

**DOI:** 10.1101/512152

**Authors:** Gray Huffman, Harrison Specht, Albert Chen, Nikolai Slavov

## Abstract

The performance of ultrasensitive LC-MS/MS methods, such as Single-Cell Proteomics by Mass Spectrometry (SCoPE-MS), depends on multiple interdependent parameters. This interdependence makes it challenging to specifically pinpoint bottlenecks in the LC-MS/MS methods and approaches for resolving them. For example, low signal at MS2 level can be due to poor LC separation, ionization, apex targeting, ion transfer, or ion detection. We sought to specifically diagnose such bottlenecks by interactively visualizing data from all levels of bottom-up LC-MS/MS analysis. Many search engines, such as MaxQuant, already provide such data, and we developed an open source platform for their interactive visualization and analysis: Data-driven Optimization of MS (DO-MS). We found that in many cases DO-MS not only specifically diagnosed bottlenecks but also enabled us to rationally optimize them. For example, we used DO-MS to diagnose poor sampling of the elution peak apex and to optimize it, which increased the efficiency of delivering ions for MS2 analysis by 370%. DO-MS is easy to install and use, and its GUI allows for interactive data subsetting and high-quality figure generation. The modular design of DO-MS facilitates customization and expansion. DO-MS is available for download from GitHub: github.com/SlavovLab/DO-MS

## Introduction

Analytical methods combining liquid chromatography and tandem mass-spectrometry (LC-MS/MS) allow for unparalleled identification and relative quantitation of the protein components of biological systems.^1–4^ Advances in LC-MS/MS have enabeled analysis of protein complexes and their functions,^5–9^ regulation of protein synthesis and alternative RNA translation,^10,11^ rare cells in blood,^12,13^ and protein conformations.^14–16^ The increasing sensitivity,^17–23^ throughput, and robustness^24^ of LC-MS/MS set the stage for quantifying thousands of proteins across many thousands of single cells, providing data with transformative potential for biomedical research.^25–29^

While LC-MS/MS proteomics methods are very powerful, they require extensive optimization of interdependent instrument parameters. Optimization is particularly critical for quantifying low-input samples, e.g., single-cell proteomes and exosomes. LC-MS/MS optimization and quality control (QC) can be performed by manually inspecting the features of peptide standards^30–32^ within MS instrument software, e.g., Thermo Scientific Xcalibur, or by specialized software packages. Following the National Institute of Standards and Technology’s QC data analysis pipeline,^33^ numerous QC programs have been developed. These programs capitalize on advances in search engines,^34^ video analysis,^35^ direct analysis of raw data,^36^ and database management tools^37^ to track instrument performance over time and assist in duty cycle optimization. Other platforms, such as MSStatsQC 2.0, employ specialized statistical methods to differentiate normal variation in instrument performance from novel variation as a means to detect instrument problems early.^38^ A comprehensive review of LC-MS/MS QC and optimization tools has been published by Bittremieux et al.^39^

We found these tools useful in developing Single-Cell Proteomics by Mass Spectrometry (SCoPE-MS), which combines TMT-labeled peptides from single cells with a TMT-labeled carrier channel to enable quantifying proteins across many single cells.^25–27,40^ However, none of these tools provided all of the metrics needed for optimizating our SCoPE-MS analysis.^40^ This motivated us to develop DO-MS, a highly modular and interactive environment for optimizing ultra-sensitive LC-MS/MS methods.

DO-MS aims to diagnose problems and suggest solutions as specifically as possible. To illustrate this point, here we describe concrete examples, including optimizing apex targeting, assessing contamination, and evaluating SCoPE-MS results. In order to enable specific diagnosis, the DOMS dashboard juxtaposes distribution plots of data from multiple levels of LC-MS/MS analysis, including retention lengths at base and mid height, intensity of all ions and of precursors selected for MS/MS, elution peak apex offset, number of MS/MS events, MS2-level coisolation (i.e., parent ion fraction), number of identified peptides at all confidence levels and quantification benchmarks. These features are organized thematically in the dashboard for ease of reference. DO-MS has already enabled us to quickly identify bottlenecks in our methods, their exact origin in the workflow, and potential solutions. Below, we share DO-MS along with a selection of examples from our work in the hope that it will facilitate the wider adoption and advancement of single-cell proteomics. We also hope that the modular nature of our platform will enable the community to add new modules for optimizing LC-MS/MS for an expanding array of applications.

## Materials and Methods

### Implementation

DO-MS is implemented as a Shiny app, built using R.^41^ All plots are generated using the ggplot2 package.^42,43^ Shiny was chosen for its interactivity, allowing data to be dynamically subset based on experiment or confidence of peptide spectral match. Additionally, this package can be run from the command line or the RStudio IDE. DO-MS is available from the Slavov Lab GitHub page: github.com/SlavovLab/DO-MS

### Data Preprocessing

The experimental data used here were generated as part of developing and optimizing Minimal ProteOmic sample Preparation (mPOP)^27^ and SCoPE-MS,^40^ and a full description of the experiments can be found in Specht et al.^27^ Briefly, all samples were separated on a 25cm length × 75*μm* Waters nanoEase column (1.7 *μm* resin, Waters PN:186008795) run by a Proxeon Easy nLC1200 UHPLC (Thermo Scientific). All samples were analyzed by a Thermo Scientific Q-Exactive mass spectrometer. Prior to running DO-MS, RAW files were searched using MaxQuant 1.6.0.16.^44–46^ The human SwissProt FASTA database (39748 entries, downloaded 5/1/2018) was used for searching data from U-937 and Jurkat cells. MaxQuant searches were conducted as previously described^27,46^. Trypsin was specified as the digest enzyme, and a maximum of two missed cleavages were allowed for peptides between 5 and 26 amino acids long. Methionine oxidation (+ 15.99491 Da) and protein n-terminus acetylation (+42.01056 Da) were specified as variable modifications. The allPeptides.txt, evidence.txt, msmsScans.txt, and msms.txt files output by MaxQuant were imported by DO-MS for analysis and figure generation. In order to make the greatest use of the DO-MS platform, users must enable the Calculate Peak Properties option on the Advanced submenu of MaxQuant’s Global Parameters tab^44,46^. This option can be automatically enabled by using the mqpar.xml files from the supporting files of this manuscript. Searches conducted without enabling this option will not generate plots for the Elution Peak Apex Offset and Peak Width at Full Width Half Max panels of the DO-MS dashboard. DO-MS is currently optimized for MaxQuant search results but users can customize it to work with alternative search-engine outputs by specifying the column headers corresponding to the data selected for visualization.

### Visualization

DO-MS’s diagnostic plots have been organized into the following five categories: Chromatography, Instrument Performance, Contamination, Peptide Identifications, SCoPE-MS Diagnostics, and DART-ID^47^. Data are visualized as full distributions using vertically oriented histograms to avoid kernel-smoothing issues. This approach is advantageous as distinct data sets may have similar summary statistics but markedly different distributions of data points.^48,49^ The full distributions allow subpopulations of ions to be identified, which can be key to optimizing LC-MS/MS performance. Additionally, these distributions can be conditioned on common ions, allowing for a more principled comparison, as discussed in the Results section.

Data imported into DO-MS can be subset based on confidence of peptide spectral match assignment and experiment name using a slider and dynamically populated list, respectively. DO-MS relies on the posterior error probability (PEP), as estimated by MaxQuant, to indicate the confidence of a given peptide spectral match (PSM). The PEP value can be thought of as the probability that the identified peptide was not in the mass spectrometer at the time the spectra was acquired.^50^ By default, DO-MS labels experiments by their corresponding raw file names. If desired, experiments can be labeled via a text-input field. Such labeling can enhance the clarity of figures and thus facilitate their analysis and broader interpretability if the figures are intended for publication.

### Report Generation and Figure Output

To facilitate the sharing of experimental results, users can output the DO-MS dashboard plots as an HTML report which reflects all data subsetting performed in the app, as well as user-supplied experimental labels. Report generation is achieved via a button in the “Report Generation” dashboard tab. Dashboard plots can also be saved as individual.png or .pdf files for use in presentations and publications directly from the dashboard tabs on which they appear.

### User Customization

We built DO-MS as a modular application to make dashboard customization and expansion as easy as possible. Each plot is generated from a separate R file, and a template file has been provided as a guide for users interested in including additional plots in their dashboard. Adding plots to the DO-MS dashboard can be accomplished by adding a customized template.r file to a new folder in the “modules” directory. After reloading the app, the new plot will automatically appear in the DO-MS dashboard in the user-specified tab. More details for this process can be found on the Github project page: github.com/SlavovLab/DO-MS

## Results and Discussion

### Sampling the Elution Peak Apex

LC-MS/MS methods aim to sample the elution profile of each peptide at its apex because such sampling maximizes the number and purity of sampled ions.^51^ However, no exisiting method can target the apex of every ion. Rather, instrument parameters can be optimized to maximize the fraction of ions sent for MS/MS at or close to their apexes. DO-MS facilitates this optimization by visualizing the distribution of apex offsets (i.e., the time offset between the apex of an ion and the time when it is MS/MSed) as estimated by MaxQuant.

The visualization of apex offsets, as shown in Fig. 1a, enabled us to rationally optimize the instrument methods for analyzing SCoPE-MS sets. To resolve the reporter ions of a TMT 11plex, we perform the MS2 scans at 70,000 resolving power, which on our Q-exactive takes 256 ms.^52^ Since Q-exactive instruments can perform the ion accumulation and MS scan in parallel, we started by setting our max fill (ion accumulation) times for MS2 scans to be 250ms.^52^ This setting resulted in sampling most ions for MS2 scans too early, significantly before the apexes of their elution peaks, Fig. 1a. We reasoned that such premature sampling could be alleviated by elongating the duty cycle either by increasing the number of MS/MSed ions per cycle or by increasing the max fill times. Indeed, increasing the fill times to 500ms and 1000ms, while keeping all other parameters constant, increased the fraction of ions whose elution peaks are sampled at or near the apex, Fig. 1a.

**Figure 1:**
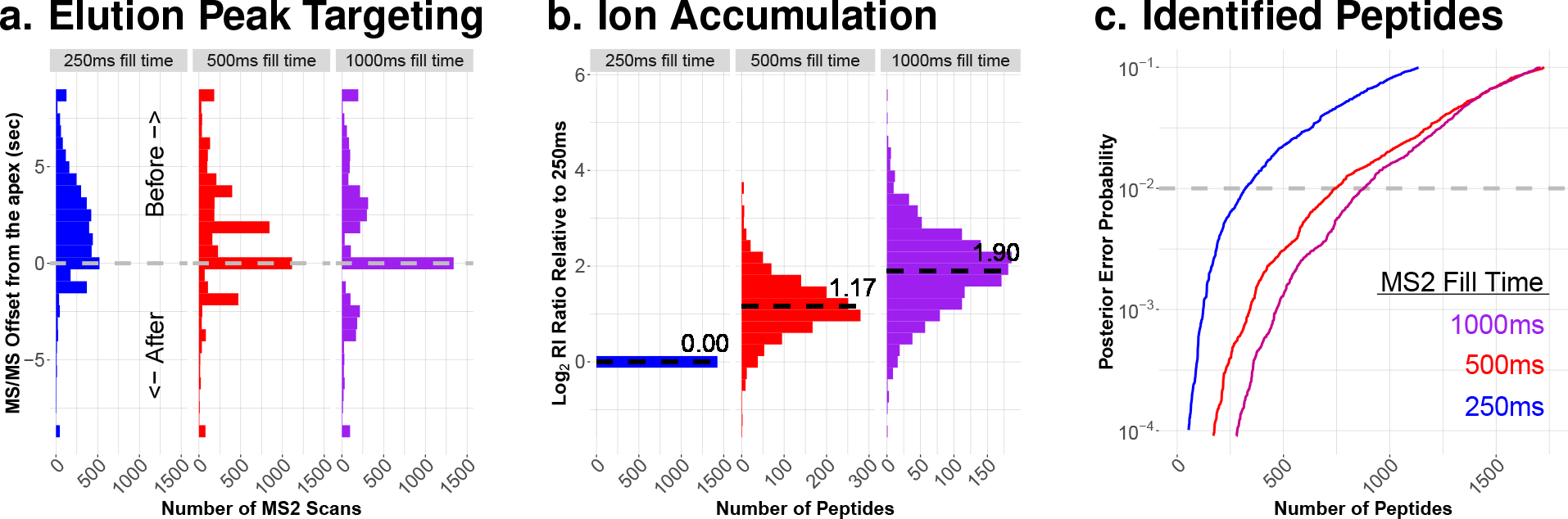
By rationally optimizing apex targeting, DO-MS increased ion delivery and peptide identification rates. (**a**) Distributions of apex offsets for three runs on 90-minute gradients. Each injection was of 1 *μl* from the same vial and corresponds to 1 × *M* dilution of a SCoPE-MS master sample as described by Specht et al.^27^ All LC parameters and instrument parameters were set to be the same except for the max fill time. (**b**) The relative efficiency of delivering ions for MS2 analysis was estimated by the intensities of reporter ions (RI). For each peptide identified across all 3 experiments, the RI intensity was divided by the corresponding RI intensity for 250ms fill time and the results for all peptides are shown as distributions on a log_2_ scale. (**c**) The number of peptides identified at each PEP threshold is shown as a rank sorted list for the three fill times. This display shows the number of peptides for all levels of confidence of identification, as quantified by the PEP.

The increased fill times and improved apex targeting increased ion delivery for MS2 analysis (Fig. 1b) and the number of identified peptides at all levels of confidence, as shown in Fig. 1c. Rather than displaying the number of identified peptides at an arbitrary confidence cut-off, DOMS plots all peptides rank-sorted by the posterior error probability (PEP) of their identification, Fig. 1c. These curves show the number of identifications for all levels of confidence. The dashed grey line on the plot denotes peptides with a 99% probability of having been correctly identified. Such curves offer insight into low-confidence peptide identifications which might be boosted by incorporating additional features, such as retention time.^46,47^

This example is consistent with previous observations that longer accumulation times can increase the number of confident peptide identifications for lowly abundant samples^17,40^. Furthermore, this example underscores that fill times can strongly influence apex targeting. Such optimization of duty-cycle and apex-targeting is sample and system dependent, and thus it requires methods that allow for rational optimization of all levels of MS analysis, such as DO-MS.

### Characterizing Contamination

Contaminants of non-protein origin are very common in proteomics experiments.^34,53^ At best, they are lowly abundant and elute separately from peptides; at worst, they are highly abundant and elute with peptides, undermining ionization, charge determination, and identification. Low-input samples are especially sensitive to contaminants, as the ratio between target and contaminant ions is more likely to be lower. DO-MS displays contaminants across the LC gradient by plotting the intensity of ions with a +1 charge state (z=1) for each minute of the gradient. This type of data presentation allows users to distinguish between hydrophilic and hydrophobic contaminants.

Due to the mosaic structure of the DO-MS dashboard, potential relationships between factors such as contaminant ion intensity and peptide identifications can be easily seen, as exemplified in Fig. 2. The juxtaposition of summed precursor intensities for contaminant ions (Fig. 2a) and for peptides (Fig. 2b) shows a clear correlation: Sample 2 has more hydrophobic contaminants and their elution coincides with reduced peptide identifications. This correlation immediately suggested that the lower identification rates for sample 2 were due to its contamination, which we subsequently identified to be polyethylene glycol (PEG).

**Figure 2:**
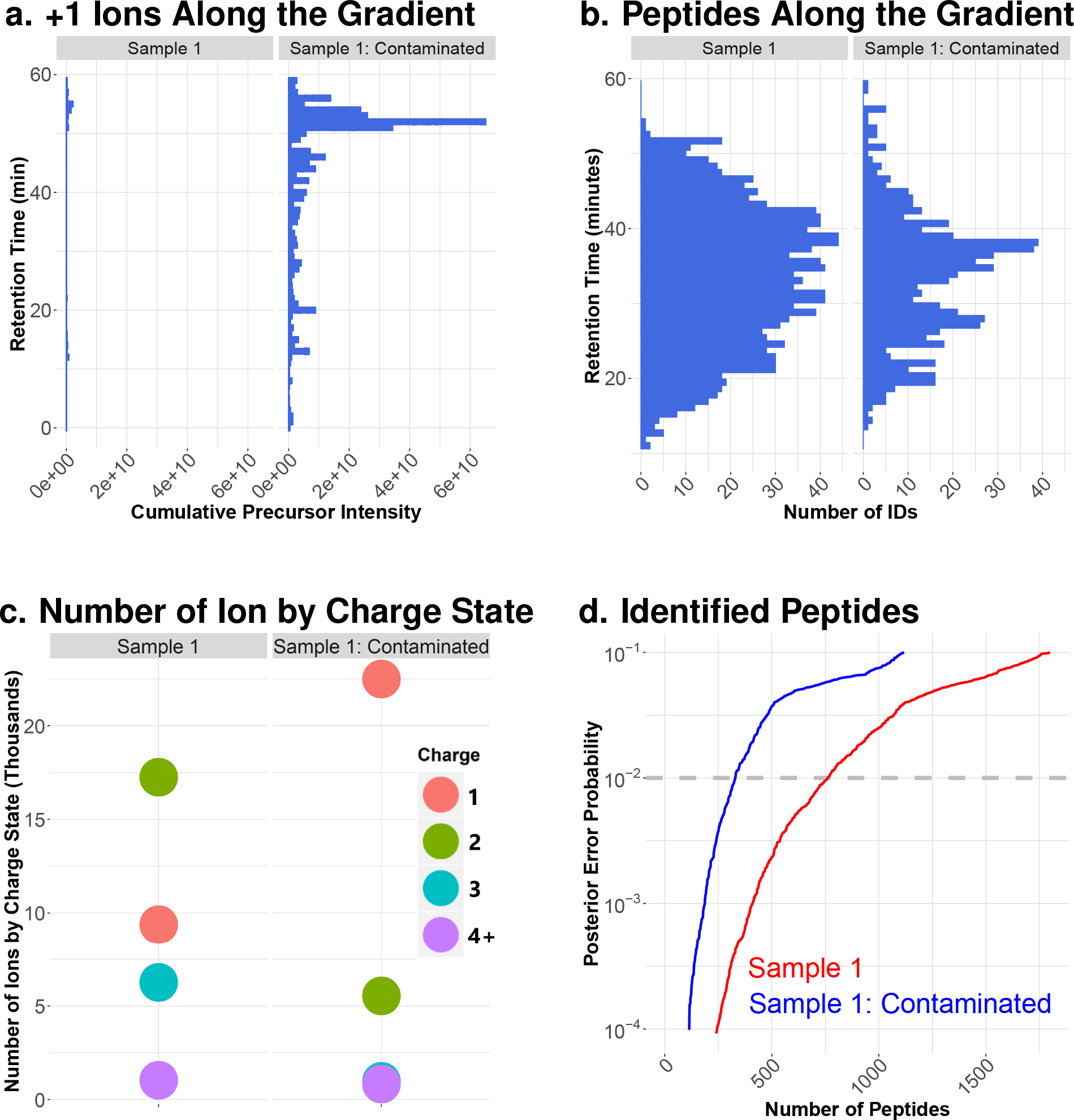
Diagnosing reduced peptide identification due to co-eluting contaminants. Plotting the cumulative intensities for all +1 ions detected during the survey scans (**a**) alongside the number of peptides identified across the gradient (**b**) can reveal correlations between co-eluting contaminants and reduced peptide identification. (**c**) Number of all detected ions by charge states. (**d**) Peptides were rank sorted by their PEPs to display the number of identified peptides across all levels of confidence.

As an additional diagnostic plot for contamination, DO-MS displays the number of ions detected by the instrument by charge state as shown in Fig. 2c. This compact display is particularly useful for comparing ions likely to be contaminants (with charge +1) and those likely to be peptides (with charge ≥ 2) across many runs. The negative impact of these numerous contaminant ions on peptide identifications can be seen in Fig. 2d. Once the presence of contaminants has been diagnosed with DO-MS, several existing software tools can be applied to more fully characterize the type of contamination present in each sample.^34,53^

### Controlled Sample Comparisons

While distributions are much more informative than their summary statistics, comparing distributions for different populations can still be misleading. Thus, when comparing the distributions plotted by DO-MS, it is important to control for (condition on) the composition of the distributions, e.g., the peptides comprising each distribution. The importance of such controlled comparison is exemplified in Fig. 3 with an experiment testing the effect of calcium addition on trypsin digestion. For this experiment, a sample was split into two equal parts, sample 1 and sample 2. They were processed in identical ways, except that 50*mM CaCl*_2_ was added to sample 2 during its digestion.

**Figure 3:**
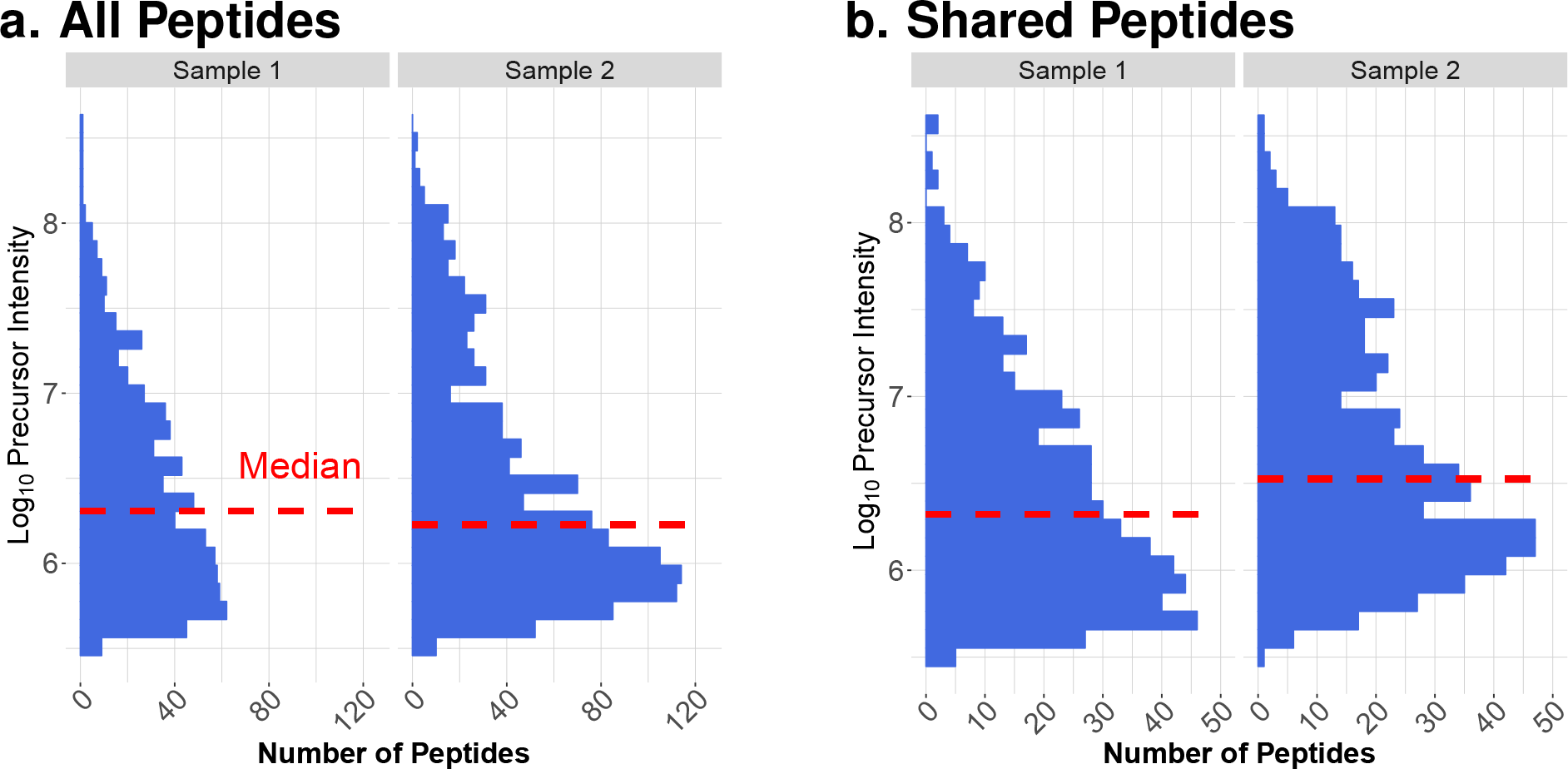
Controlled comparison of peptide abundances across experiments. Without controlling for the composition of the two populations being compared, trends in the data can be misread. In this case, when comparing the distribution of precursor intensities for all peptides identified in each sample (**a**), Sample 1 appears to have more highly abundant peptides. However, when ensuring that the comparison is only based on those peptides identified in each sample (**b**), the opposite trend becomes apparent: namely, that the peptide species in Sample 2 were more highly abundant.

Both samples were digested with 20*ng/μl* Promega Trypsin/ LysC mix. The distributions of all peptide abundances and the corresponding median abundances (Fig. 3a) may be interpreted to suggest that sample 1 resulted in more efficient delivery of peptides to MS analysis and the addition of calcium was detrimental. However, conditioning the comparison on the common peptides indicates the opposite conclusion: The addition of calcium chloride to sample 2 during its digestion resulted in more efficient delivery of peptides to the instrument.

### Low-Input Sample Diagnostics

In the process of developing Minimal ProteOmic sample Preparation (mPOP)^27^ and SCoPE-MS,^40^ we discovered a number of helpful metrics for diagnosing SCoPE-MS sample preparation and optimizing instrument parameters for low-input samples. Two such metrics are relative reporter ion (RI) intensities and the correlations among them. These metrics indicate the efficiency of single-cell preparations and quantitative data quality.

To optimize SCoPE-MS, we used bulk SCoPE-MS sets from which a single injection (1%) corresponds to a single-cell SCoPE-MS set. Such samples allowed us to assess and optimize instrument performance independent of sample variability (since we could inject multiple aliquots of the same sample) and to assess quantification, as we had a strong expectation that the pseudo-single-cell channels should correlate positively to their corresponding carrier channel. Below we demonstrate diagnosis of such a sample by DO-MS; the sample preparation is described in detail by Specht et al^27^. Briefly, the sample was composed of 10 TMT channels containing serial dilutions of digested cell lysate from two cell types: U-937 (monocytes, denoted by U) and Jurkat (T-cells, denoted by J). This reference standard is diluted so that 1 *μl* corresponds to a SCoPE-MS set and contains peptide input equivalent to about 106 single cells (20 — 50*ng* of total protein). The reference samples have two carrier channels, each of which contains peptide input comparable to 50 cells of each type, and six channels contain peptide inputs comparable to individual single cells, three of each type.^27^

The distribution of reporter ion (RI) intensities from a SCoPE-MS set can be an informative indicator. Low RI intensities may be due to failed cell isolation, digestion, or labeling. High RI intensities may be due to background contamination or cross-labeling. Such deviations may be diagnosed simply from the distributions of RI intensities. However these distributions are quite broad since the RI intensities for quantified peptides often span several orders of magnitude. To decrease this dynamic range, DO-MS plots the distribution of relative RI intensities: for each peptide, the RI intensities are scaled by dividing them by the RI intensity in the most abundant channel (the one with the highest median RI intensity), Fig. 4a. This visualization clearly indicates whether the single-cell channels have about 50-fold lower median intensity than a 50-cell carrier channel, as well as the background signal (including isotopic contamination) in the empty channels. Such a diagnostic is helpful for determining label-quenching efficiency, as well as the efficiency of cell sorting by FACS. By examining the relative RI intensities present in blank channels, one can also assess the amount of background signal present and the degree of isotopic carryover.

**Figure 4:**
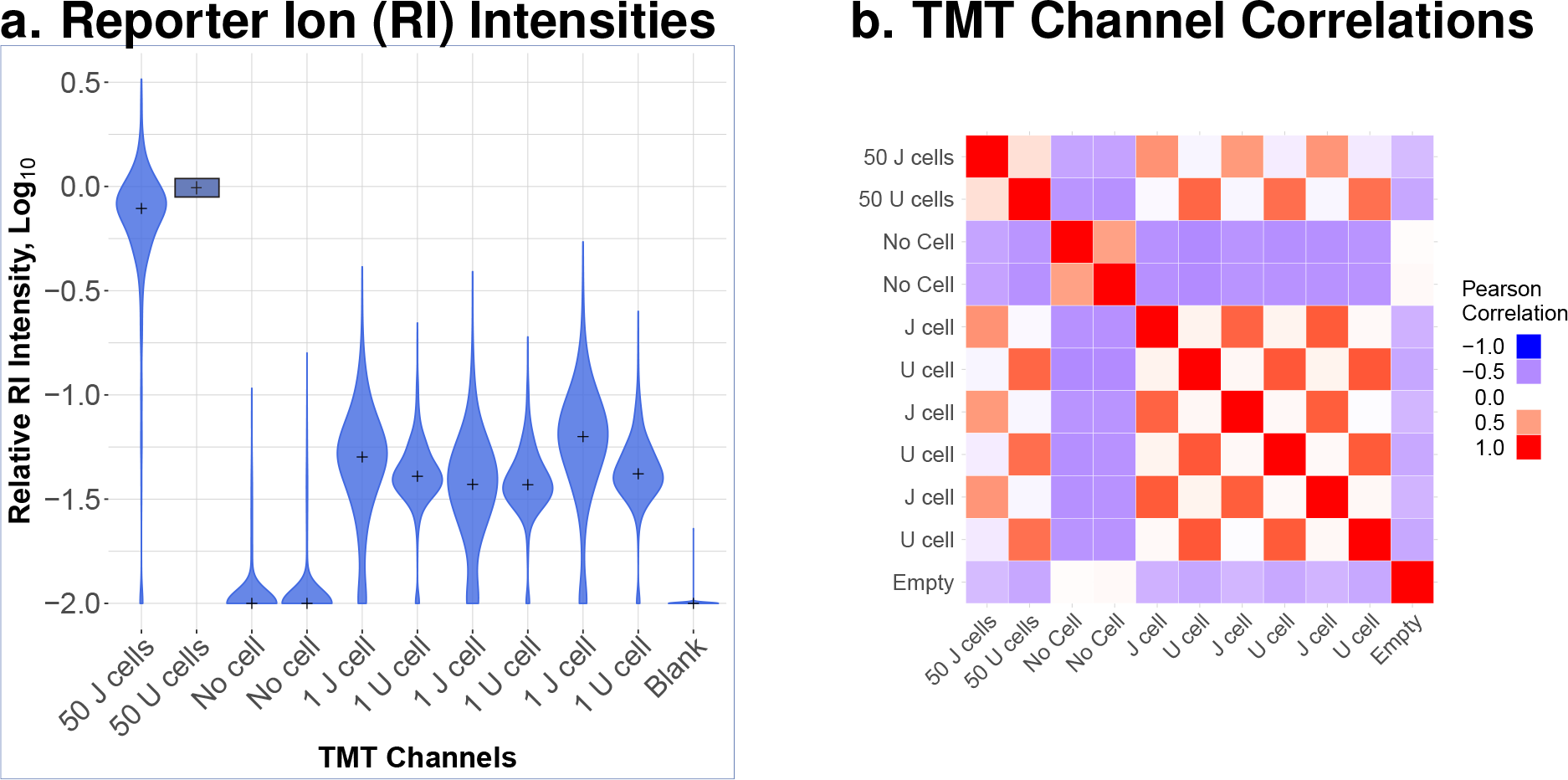
Evaluating low-input samples, such as SCoPE-MS sets. (**a**) the distributions of relative reporter ion (RI) intensities can indicate the relative amount of peptides and the efficiency of sample preparation for each channel. (**b**) the matrix of pairwise correlations among all all channels of a SCoPE-MS set can be used to benchmark relative quantification within that set. In (**a**) we expect single-cell channels to have relative RI intensities that are 50-fold lower than the 50-cell carrier channels (about 1.5 on log_10_ scale). In (**b**), we expect singlecell channels to correlate positively with single-cell channels and carrier channels that contain their respective cell type, while cross-cell-type correlations for single-cell channels are expected to be negative.

The quantitative accuracy of SCoPE-MS sets can be benchmarked by comparing the relative quantification from the carrier channels and single-cell channels as quantified by the correlations among them. DO-MS computes all possible pairwise correlations, i.e. the correlation matrix for the column and row-normalized RI intensities. This matrix can serve as a complimentary diagnostic for relative quantification in SCoPE-MS sets based on the expectation that cells of the same type should correlate positively with each other but not with different cell types. Furthermore, blank channels should not correlate positively with either the single-cell or carrier channels. This expectation is consistent with the correlations shown in Fig. 4b. This correlation matrix can also be useful for identifying cross-contamination, cross-labeling, and on-column carryover.

These SCoPE-MS plots should be analyzed in the context of distribution plots reporting on all levels of the LC-MS/MS analysis so that the origin of problems can be identified and interdependent parameters optimized. We hope that these metrics will assist the wide adoption of low-input sample preparation and analysis methods.

## Conclusion

Optimal instrument parameters are context dependent and thus should be determined systematically by data-driven approaches for each set of samples and LC-MS/MS configurations. DO-MS enables such optimization. Sometimes, this data-driven approach results in counter-intuitive results, as demonstrated in Fig. 1 with the increased number of identified peptides at longer max fill times. By increasing our MS2 injection time, and consequently our duty cycle length, we managed to increase the number of confidently identified peptides in our sample. This result contrasts with the strategy commonly employed by bulk proteomics methods, namely seeking to increase peptide identification by increasing MS2 sampling frequency and thus decreasing the fill time for each MS/MS.

The DO-MS dashboard is an open-source, GUI-based tool for quickly assessing LC-MS/MS parameter optimization strategies and sample quality in single-cell proteomics experiments. This diagnostic platform can assist other labs in adopting ultrasensitive, low-input LC-MS/MS methods, such as SCoPE-MS, and can serve as a highly adaptable data visualization tool for proteomics researchers.

## Availability and Documentation

The DO-MS Shiny dashboard, basic documentation, and a sample data set are currently available through a git repository at github.com/SlavovLab/DO-MS, under the MIT license. A free, web-based version of the DO-MS is also accessible at singlecellproteomics.shinyapps.io/QCtest/, although it is recommended that interested users install the dashboard locally to avoid data upload restrictions and monthly activity caps.

## Acknowledgments

The catalyst for creating this package was a break-out discussion group led by Jürgen Cox at the 2018 Single-Cell Proteomics conference. Special thanks goes to Jürgen Cox for structuring this discussion, as well as to Ezra Levy, Guillaume Harmange, and Hung-Yi Wu for their part in outlining the characteristics that an instrument-parameter optimization and sample-quality analysis software package should possess. We would also like to thank Ed Emmott, David Perlman, and Toni Koller for their valuable feedback and troubleshooting efforts during the package development process.

